# Membrane Expression Enhances Folding, Multimeric Structure Formation, and Immunogenicity of Viral Capsid Proteins

**DOI:** 10.1101/2024.10.06.616851

**Authors:** Junru Cui, Fangfeng Yuan, Jane Qin, Ju Hyeong Jeon, Dong Soo Yun, Tianlei Wang, Renhuan Xu, Helen Cao, Jianzhu Chen

## Abstract

Viral capsid proteins are widely explored for subunit vaccine development but are often hampered by their complexity of production and low immunogenicity. Here, we report a simple approach to overcome these challenges by combining mRNA vaccine technology with protein engineering. Using African swine fever virus (ASFV) capsid proteins P72 and penton as models, we engineered them into membrane-bound and secreted forms, and compared their immunogenicity to the native intracellular form in mice and pigs through mRNA vaccination. The membrane-bound and secreted P72 and penton folded into their native multimeric structure independent of viral chaperone, therefore preserving their conformational epitopes. The membrane-bound P72 and penton also elicited significantly stronger antibody and T cell responses than their secreted or intracellular counterparts. Our study provides a simple approach to enhance folding, multimeric structure formation, and immunogenicity of viral capsid proteins for ASFV subunit vaccine development and immunogenicity of intracellular proteins in general.

## Main

Vaccination remains a cornerstone of infectious disease control. Traditional vaccines based on inactivated and live-attenuated infectious agents have provided protection against many pathogens, but they are either ineffective or have unacceptable safety concerns in protecting against certain pathogens. For example, African swine fever virus (ASFV) causes a highly contagious and deadly disease in both domestic and wild swine, with devastating economic consequences for the global pork industry^1–3^. Inactivated ASFV vaccines do not confer protection while the live-attenuated vaccines are effective but with high risk of reversion to virulence^4,5^. Subunit vaccines based on recombinant proteins and nucleic acids offer greater precision and safety^6^. Central to the success of the subunit vaccine development is the design of antigens capable of eliciting strong and protective immune responses.

Viral envelope and capsid proteins are among the most explored antigens for subunit vaccine development. Naturally, envelope proteins are displayed on the surface of virion (or infected host cells) in large numbers. The resulting high epitope density and accessibility to B cell recognition induces potent antibody responses. In contrast, viral capsid proteins are synthesized in the cytosol and often require chaperone for folding and assembly into capsid for packaging viral genome. The use of capsid proteins in subunit vaccine is often hampered by the complexity in their production due to susceptibility to misfolding and dependence on specific post-translational modifications, and poor immunogenicity as recombinant capsid proteins may not fold into their natural multimeric structures for inducing antibody responses against conformational epitopes that many neutralizing antibodies target^7^. A more successful approach has been to express and assemble viral capsid proteins into virus-like particle (VLP), as demonstrated by VLP-based vaccines for hepatitis B (HBV)^8^, human papillomavirus (HPV)^9^, and hepatitis E (HEV)^10^. Although VLP preserves the conformational epitopes and the high epitope density of the natural viral capsids and therefore induces potent antibody responses, it does not induce any significant CD8^+^ T cell response. In additional, VLP production in scale has not been possible for many pathogenic viruses due to the complexity of capsid assembly.

Here, we report a novel approach to harness capsid proteins for subunit vaccine development by preserving their high-density conformational epitopes and their ability to induce both humoral and cellular immunities without the technical challenges of VLP assembly. We used the ASFV capsid proteins as a proof-of-concept by combining mRNA vaccine technology and advanced protein engineering. The outer shell of ASFV capsid is composed of the major capsid protein P72, which exists as a homotrimer in the capsid, and the minor capsid protein penton, which exists as a homopentamer in the capsid^11,12^. Studies have suggested P72 as one of the most promising targets for ASFV subunit vaccine development (Reviewed in^13–15)^. However, previous attempts to utilize recombinant P72 in subunit vaccine development have been hindered by its tendency to form monomers rather than the more immunogenic native trimers^16–19^ and the requirement for chaperone pB602L for folding^20,21^. In comparison, the immunogenicity of penton has never been explored^11,12^. Specifically, we engineered both P72 and penton into membrane-bound and secreted proteins and compared their immunogenicity to the native intracellular form in mice and pigs through LNP mRNA vaccination. Notably, the membrane-bound and secreted P72 were able to fold into trimers independent of chaperone pB602L and the membrane-bound and secreted penton also form native pentamer, therefore preserving their conformational epitopes. Immunogenicity testing in both mouse and pig models showed that the membrane-bound P72 and penton elicited significantly stronger antibody and T cell responses than their secreted or intracellular counterparts. Our study demonstrates a proof-of-concept for enhancing folding, formation of native multimeric structures, and immunogenicity of viral capsid proteins, offering a simple strategy for developing safe and effective subunit vaccines for ASFV. More broadly, the same approach should improve immunogenicity of any intracellular proteins of pathogen origin or otherwise for preventive and therapeutic vaccination.

## Results

### Folding and trimer formation of the membrane-bound and secreted P72 do not require chaperone

Recent structural studies revealed that the outer capsid shell of ASFV is composed of a major capsid protein P72, which forms homotrimer, and a minor capsid protein penton, which forms a homopentamer^11,12^. The outer capsid shell is stabilized by three minor proteins underneath, including P17 that glues together P72 trimers^11,12^. To enhance immunogenicity of P72 and penton by maintaining their native multimeric structures and therefore conformational epitopes, we expressed P72 and penton as the membrane-bound (MB) and secreted (S) proteins, in additional to their native intracellular (IC) form. Specifically, to express P72 as a secreted protein (S-P72) we added the signal peptide of human CD8α to the N-terminus of P72 (Figure 1A). To express P72 as a membrane-bound protein (MB-P72), in addition to the signal peptide, we added the hinge domain (HD), transmembrane domain (TMD), and a short cytoplasmic region from the same CD8α to the C-terminus of P72. CD8α signal peptide, hinge and transmembrane domains were chosen because they are widely used to express chimeric antigen receptors on the cell surface^22^. AlphaFold2^23^ and PyMOL modeling suggested that the addition of the CD8α hinge domain does not induce significant structural alterations of P72 nor affects the formation of P72 trimer probably because the long CD8α hinge domain provides sufficient flexibility and space to accommodate P72 trimer formation (Extended Fig. 1A,C), consistent with a previous report^24^.

**Figure 1.**
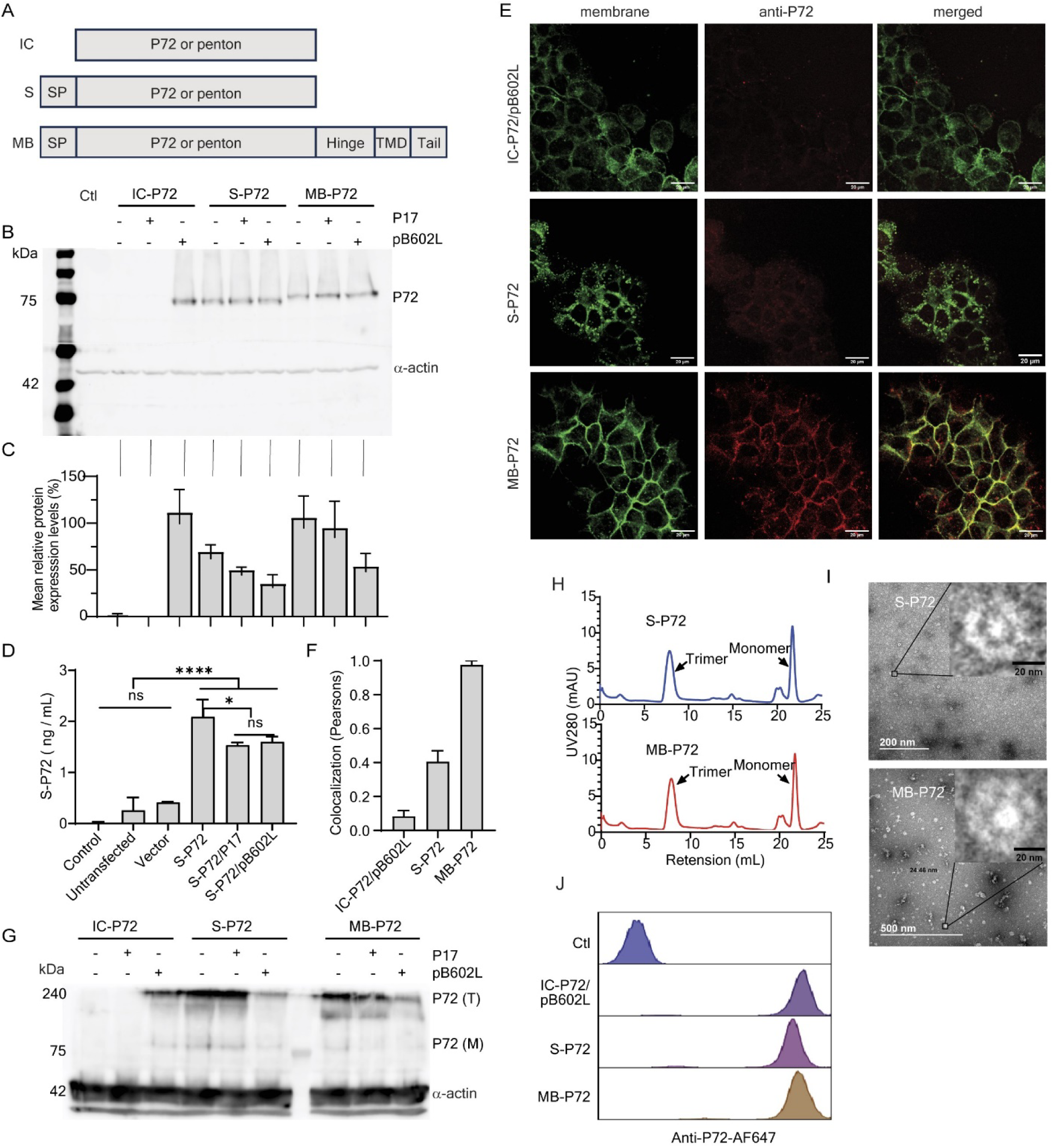
Characterization of three forms of P72 and their trimer formation. **(A)** Schematics of intracellular (IC), secreted (S) and membrane-bound (MB) P72 or penton. Secreted P72 or penton contains a signal peptide (SP) at the N-terminus. In addition to SP, the membrane-bound P72 or penton contains the CD8α hinge, transmembrane domain (TMD), and a truncated cytoplasmic tail. **(B)** A representative Western blot showing expression of IC-P72, S-P72 and MB-P72 alone or in the presence of P17 or pB602L. HEK293T cells were transfected with IC-, S- and MB-P72 alone or together with P17 or pB602L. Cell lysates were subjected to Western blotting analysis 48 hours later and probed with anti-P72 and anti-α-actin. **(C)** Relative expression levels of IC-, S- and MB-P72 normalized to IC-P72/pB602L. Results are mean ± standard deviation (SD) based on four independent experiments in HEK293T cells and Vero cells. **(D)** Detection of S-P72 in culture supernatants by ELISA. Culture supernatants were collected from untransfected, vector transfected, and S-P72 (w/o P17 or pB602L)-transfected HEK293T cells, followed by ELISA. P72 concentrations were determined through a standard curve and presented as the mean ± SD based on three technical replicates. Control was without culture supernatant. Statistical analysis was conducted using one-way analysis of variance (ANOVA) against empty vector transfected cells. * p<0.05 and ****p<0.0001. **(E)** Representative images showing co-localization of P72 (red) and the cell membrane (green) in Vero cells transfected with IC-P72/pB602L, S-P72, and MB-P72 for 48 h. IFA was performed using a primary mouse anti-P72 monoclonal antibody followed by a secondary goat anti-mouse IgG conjugated with AF647. Cell membrane was stained with CellMask™ Plasma Membrane Stains (green). **(F)** Pearson correlation coefficient of IC-, S- and MB-P72 and cell membrane as measured by ImageJ in two separate experiments as shown in E. **(G)** A representative Western blot showing trimer formation following expression of IC-P72, S-P72 and MB-P72 alone or in the presence of P17 or pB602L. HEK293T cells were transfected with IC-, S- and MB-P72 alone or with P17 or pB602L. Cell lysates were subjected to SDS-PAGE under nonreducing condition followed by Western blotting with anti-P72 and anti-α-actin. The P72 monomer and trimer are indicated as P72 (M) and P72 (T), respectively. **(H)** Size exclusion chromatography analysis of purified HA-tagged S-P72 (top panel) and MB-P72 (bottom panel) samples. P72 trimer and monomer are labeled. **(I)** Transmission electron microscopy visualization of purified HA-tagged S-P72 and MB-P72 by negative staining. Inserts show representative enlarged top view images of S-P72 and MB-P72. Scale bars are shown. **(J)** Flow cytometry analysis of P72 expression following transfection of HEK293T cells with LNP-mRNA encoding IC-, S- and MB-P72. Cells were permeabilized and stained with mouse anti-P72 followed by AF647-labled goat anti-mouse IgG. Control (Ctl) was untransfected cells stained the same way.

We determined the requirement of chaperone pB602L, a viral protein known to facilitate P72 folding and trimer formation^20,21^, and P17 in P72 expression. HEK293T and Vero cells were transiently transfected with vectors expressing IC-, S- and MB-P72 with or without co-transfection of P17 or pB602L, followed by Western blotting analysis of cell lysates. The level of IC-P72 expression was minimal by itself or in the presence of P17, but greatly elevated in the presence of pB602L (Figure 1B, C). In contrast, the expression of S-P72 and MB-P72 was readily detected and their level of expression was not significantly affected by co-expression of P17 or pB602L. Approximately 2 ng/mL of S-P72 was detected by ELISA in the supernatants of S-P72-transfected HEK293T cells (Figure 1D). Notably, this level was not affected by the co-expression of P17 or pB602L. Immunofluorescence assay (IFA) showed co-localization of MB-P72 with the cell membrane (Figure 1E). Based on Pearson correlation, colocalization of P72 and cell membrane was 97.5% for MB-P72, 40.5% for S-P72, and 8.3% for IC-P72/pB602L (Figure 1F).

To assay P72 trimer formation, we used three approaches. First, cell lysates were subjected to SDS-PAGE under non-reducing conditions followed by Western blotting. A dominant P72-reactive band was detected at approximately 240 kDa (Figure 1G), suggesting trimer formation. Consistently, formation of P72 trimer from IC-P72 was only detected in the presence of pB602L, while formation of P72 trimer from S-P72 and MB-P72 did not require pB602L. Second, we expressed and purified HA-tagged S-P72 and MB-P72 and quantified P72 trimers and monomers by size exclusion chromatography. As shown by Figure 1H Extended Table 2, 50.8% of S-P72 and 36.8% MB-P72 were trimers. Third, we confirmed the structural similarity of trimers formed by HA-tagged S-P72 and MB-P72 by transmission electron microscopy (TEM) (Figure 1I) to those of native ASFV virions (PDB ID: 6L2T, EMDB ID: 0814)^11^ and previously reported P72 structures (PDB ID: 6KU9, EMDB ID: 0776)^25^.

Together, these results show that while folding and trimer formation of native P72 (IC-P72) requires chaperone pB602L, folding and trimer formation of S-P72 and MB-P72 do not require pB602L, probably due to re-direction of P72 synthesis into the endoplasmic reticulum (ER) by the signal peptide.

To develop P72 mRNA vaccines, we cloned MB-P72, S-P72, IC-P72, and pB602L into the pUC57 vector, which includes a T7 promoter, 5’UTR, 3’UTR, and polyA tail. The mRNAs were synthesized via in vitro transcription using T7 polymerase^26^ and transfected into HEK293T or Vero cells using MessengerMax^TM^. P72 expression was confirmed by Western blotting analysis with relative expression of IC-P72:S-P72:MB-P72 at 1:2:8 (Extended Fig. 2). The MB-P72 and S-P72 mRNAs were formulated into lipid nanoparticles (LNP) individually and IC-P72 and pB602L mRNAs were formulated together into LNP. The resulting LNP-mRNAs exhibited a narrow size distribution of approximately 115 nm and an encapsulation efficiency exceeding 91% (Extended Table 3). Transfection of LNP-mRNAs into HEK293T cells led to robust expression of MB-P72, S-P72, and IC-P72 as detected by flow cytometry (Figure 1J), suggesting successful expression of the three forms of P72 via mRNAs.

**Figure 2.**
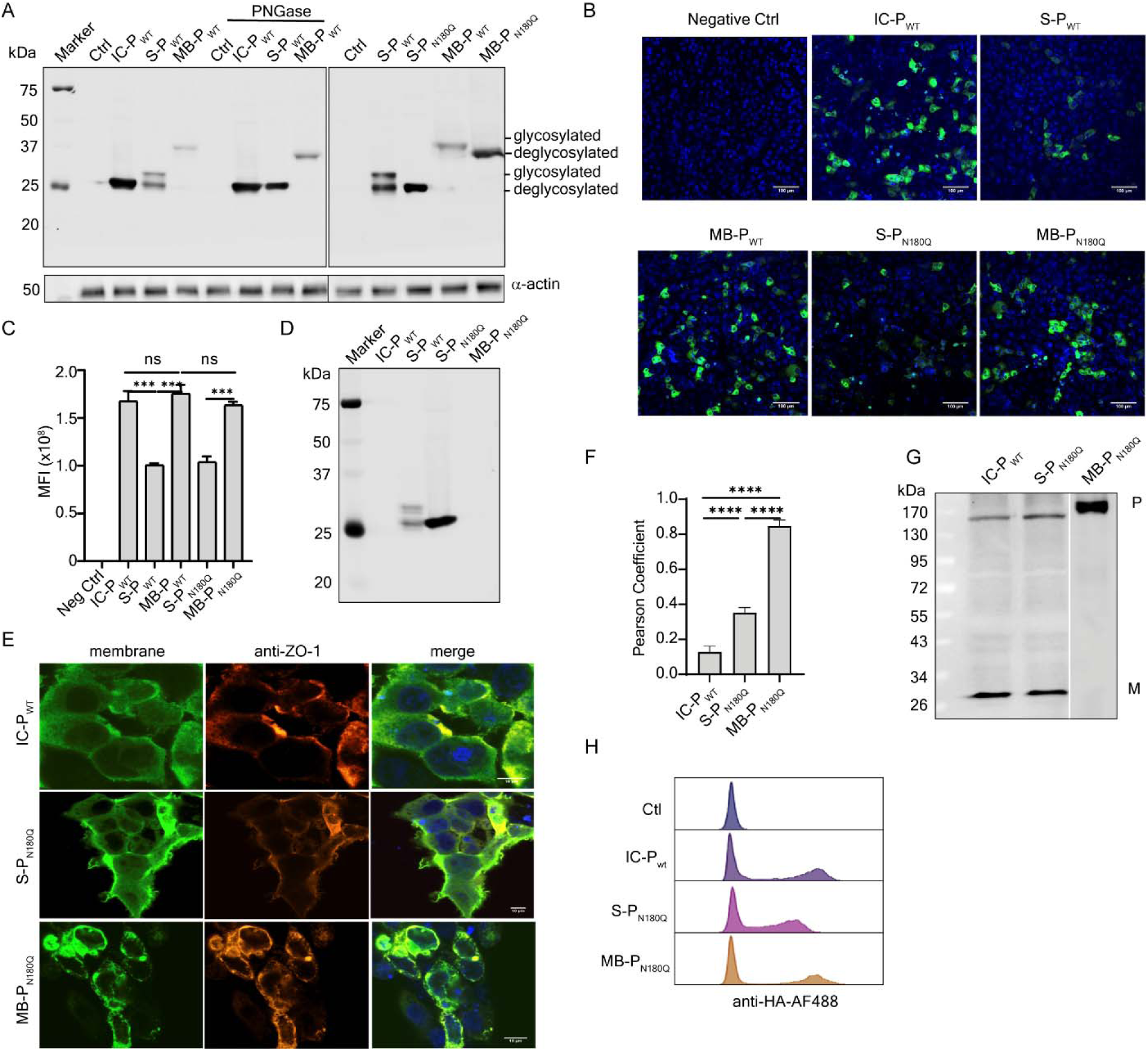
Characterization and de-glycosylation of engineered penton. (**A**) A representative Western blotting analysis of IC-, S- and MB-P_WT_. HEK293T cells were transfected with HA-tagged penton plasmids and cell lysates were prepared 48 h post transfection. Cell lysates with and without PNGase treatment were subject to SDS-PAGE followed by Western blotting using mouse anti-HA monoclonal antibody and goat anti-mouse IgG secondary antibody. P_WT_, wildtype penton, P_N180Q_, penton with N to Q mutation at position 180. (**B**) Representative immunofluorescent images of IC-, S- and MB-P_WT_ transfected HEK293T cells. (**C**) Comparison of mean fluorescence intensity (MFI) from three independent experiments. **(D)** Detection of secreted penton in cell culture supernatants. Suspension HEK293F cells were transfected with HA-tagged penton plasmids and cell culture supernatant was collected, concentrated using a 10 kDa centrifugal filter followed by Western blotting analysis as in A. (**E-F**) Representative confocal images of penton colocalization with cell membrane marker (ZO-1) (**E**) and Pearson Coefficients for colocalization of IC-, S- and MB-P_WT_ with the cell membrane (**F**). Vero cells were transfected with plasmids expressing HA-tagged penton proteins. Cells were fixed 48 hour later, permeabilized, and stained with FITC-conjugated anti-HA antibody and rabbit polyclonal anti-ZO-1 followed by Alexa Fluor™ 594-labeled goat anti-rabbit IgG secondary antibody. Experiments were performed in triplicates and repeated at least two times. Error bars in F represents mean ± SD. Statistical analysis was performed using one-way ANOVA. *** p<0.001; **** p<0.0001. **(G)** A representative Western blotting analysis of monomeric and pentameric pentons. The experiment was done in the same way as in A except that non-reducing SDS-PAGE was used. P, pentamer; M, monomer. (**H)** Flow cytometry analysis of penton expression following transfection of HEK293T cells with LNP-mRNA encoding IC-P_WT_, S-P_N180Q_ and MB-P_N180Q_. Cells were permeabilized and stained with AF488 conjugated mouse anti-HA. Control (Ctl) was untransfected cells stained the same way.

### Penton forms pentameric structure without viral chaperone

We also expressed penton, referred to as P_WT_, in membrane-bound (MB), secreted (S), and native intracellular (IC) forms (Figure 1A). Because of a lack of antibodies specific for penton, we added a HA-tag at the C-terminal of penton for easy detection. Structural modeling with PyMOL indicated that the hinge domain provides sufficient flexibility to accommodate pentamer formation (Extended Fig. 1D), consistent with a previous report^24^. HEK293T cells were transiently transfected with vectors expressing IC-, S- and MB-P_WT_ followed by Western blotting analysis of cell lysates. The IC-P_WT_ migrated at the predicted size of 28.9 kDa. S-P_WT_ produced an additional band slightly larger than that of IC-P_WT_; and MB-P_WT_ migrated slightly larger than the predicted 36.9 kDa (Figure 2A). Treatment of cell lysates with PNGase F, which removes N-linked oligosaccharides, did not affect the size of IC-P_WT_, but reduced the larger S-P_WT_ to the same size as IC-P_WT_ and the size of MB-P_WT_ slightly, indicating glycosylation of both S-P_WT_ and MB-P_WT_. Since N-linked glycosylation can shield B cell epitopes and hinder immune responses to the native protein, we identified a single NLTT glycosylation motif in the wildtype penton that is exposed and mutated N at residue 180 to Q (N180Q) (Extended Fig. 1B). S-P_N180Q_ and MB-P_N180Q_ produced a single band at the same size as the non-glycosylated counterparts (Figure 2A), suggesting successful abolition of N-linked glycosylation. Further characterization using confocal microscopy revealed that IC-P_WT_ and MB-P_WT_ were expressed at similar levels, significantly higher than S-P_WT_ (Figure 2B, C). The N180Q mutation did not significantly alter their levels of expression.

To detect the secretion of penton, culture supernatants from transfected HEK293F cells were concentrated and analyzed by Western blotting. Both non-glycosylated and glycosylated S-P_WT_ were detected in the supernatants of S-P_WT_-transfected cells, while only non-glycosylated S-P_N180Q_ was detected in the supernatants of S-P_N180Q_-transfected cells (Figure 2D). As expected, IC-P_WT_ and MB-P_N180Q_ were not detected in the culture supernatants. To investigate subcellular localization, HEK293T cells transfected with HA-tagged penton were permeabilized and stained with anti-HA and Zonula Occludens (ZO)-1, a cell membrane marker, followed by confocal imaging. MB-P_N180Q_ predominantly colocalized with ZO-1 on the cell membrane, whereas IC-P_WT_ and S-P_N180Q_ primarily localized within the cytosol (Figure 2E). Quantification of the Pearson colocalization coefficient showed that more than 80% of MB-P_N180Q_ colocalized with ZO-1, compared to only 35% for S-P_N180Q_ and 10% for IC-P_WT_ (Figure 2F). Furthermore, we directly assessed pentameric penton formation by Western blotting analysis using SDS-PAGE under non-reducing conditions. Cell lysates from IC-P_WT_ and S-P_N180Q_ transfected HEK293T cells gave rise to two predominant bands corresponding to monomeric and pentameric forms of penton (Figure 2G), whereas only pentamer was detected in MB-P_N180Q_-transfected cells. Additionally, we prepared IC-P_WT_, S-P_N180Q_, and MB-P_N180Q_ mRNAs and formulated them into LNP. Physicochemical analysis confirmed a narrow particle size distribution (∼120 nm) with over 94% encapsulation efficiency (Extended Table 3). Expression of IC-P_WT_, S-P_N180Q_, and MB-P_N180Q_ was validated by flow cytometry analysis of LNP-mRNA transfected HEK293T cells (Figure 2H).

Together, these results show that penton can form pentamer without viral chaperone and its expression in membrane-bound form further promote pentamer formation. Redirection of penton synthesis in the ER and Golgi leads to N-glycosylation, which is abolished by the N180Q mutation.

### MB-P72 is more immunogenic than S-P72 and IC-P72 in mice

To compare the immunogenicity of MB-, S- and IC-P72, we immunized BALB/c mice intramuscularly with 5 μg of mRNA in LNP formulation (using ARV L002 ionizable lipid)^26^ at day 0 and 21 (Figure 3A). Sera were collected before immunization and two weeks after each immunization (days 14 and 35) and used to assay P72-specific IgG by ELISA. Spleen and bone marrow were harvested at day 35 and used for assaying IFN-γ- and TNF-α-secreting T cells and antibody secreting B cells (ASC) by ELISPOT, respectively. P72-specific IgG titers 14 days following the first immunization were 62.4 for IC-P72 plus pB602L (shown as IC-P72* on Figs), 129 for S-P72 and 330 for MB-P72 (Figure 3B). Following the boost, the titers increased to 93 for IC-P72*, 4576 for S-P72 and 50820 for MB-P72. Consistently, the frequency of ASCs in the bone marrow at day 35 tended to be higher in MB- and S-P72 immunized mice (Figure 3C). Furthermore, the frequencies of IFN-γ- or TNF-α-secreting cells following stimulation of splenocytes with a P72 peptide pool (Extended Table 1) tended to be higher in MB-P72 immunized mice (Figure 3D, E). In a separate experiment where P72 mRNAs were formulated in LNP using ionizable lipid SM-102, MB-P72 also induced higher antigen-specific IgG responses than IC-P72* (Extended Figure 3B). These results show that MB-P72 is more immunogenic than S-P72 and IC-P72 in inducing antibody and T cell responses in mice.

**Figure 3.**
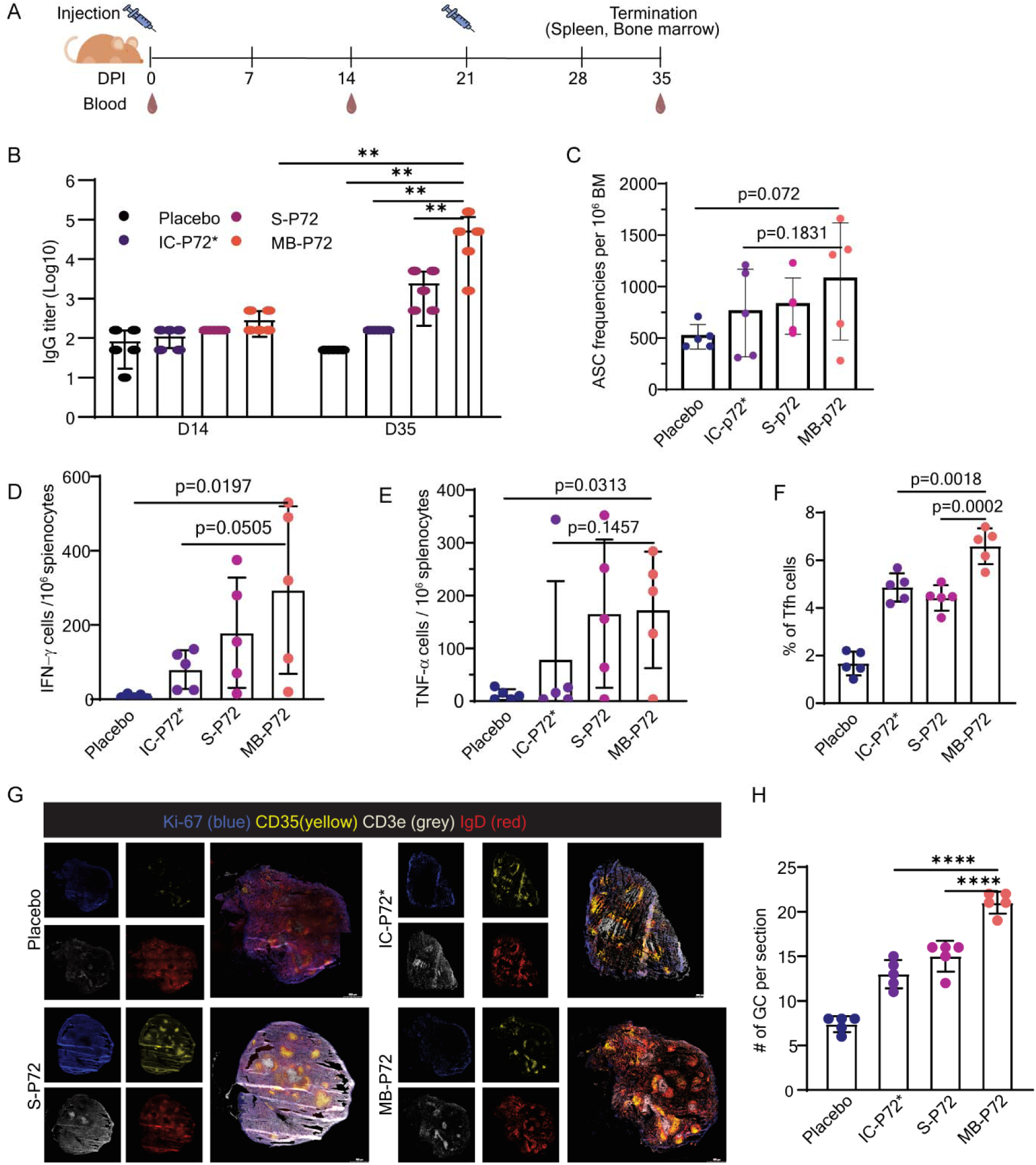
MB-P72 is more immunogenic than S-P72 and IC-P72 in mice. (**A**) Scheme of immunization. BALB/c mice (n=5 mice per group) were intramuscularly (i.m.) injected with LNP mRNA encoding IC-P72 plus pB602L (IC-P72*), S-P72 and MB-P72 and boosted at day 21. Sera were collected before and 14 days after each immunization. Spleen and bone marrow were harvested at day 35. (**B**) P72-specific IgG titers at day 14 and day 35. Placebo (blue), IC-P72* (purple), S-P72 (magenta), and MB-P72 (orange). Sera from different groups were 3-fold serially diluted and assayed for P72-specific IgG by ELISA. The titer was determined by interpolating cutoff value (OD_450_ = Average of placebo + 3 SD) from the standard curve of serially diluted samples. (**C**) The frequencies of IgG secreting cells per million bone marrow cells by ELISPOT. 10^5^ bone marrow cells were incubated in wells precoated with anti-IgG for 36 hours and IgG was detected with anti-mouse IgG detecting antibody. Antibody secreting B cells (ASC) were detected by ELISPOT reader. (**D, E**) The number of IFN-γ (D) and TNF-α (E) secreting cells (SFC) per 10^6^ splenocytes from different groups. T cells specific for P72 were assayed using single color IFN-γ or TNF-α ELISpot kit following stimulation of splenocytes with a p72 peptide pool for 36 hours. (**F**) Comparison of percentages of CD4^+^B220^-^CD44^+^PD-1^+^CXCR5^+^ Tfh cells in the spleen of immunized mice. Splenocytes were stained with antibodies specific for CD4, B220, CD44, PD-1, and CXCR5 followed by flow cytometry. Tfh cells were gated on CXCR5^+^ and PD-1^+^ population among the CD44^+^CD4^+^B220^-^ T cells (Extended Fig. 4). (**G, H**) Representative immunofluorescent images of spleen sections of immunized mice (G) and quantification of the numbers of germinal centers (H). Spleen sections were stained for IgD, CD35, CD3ε, and Ki-67, imaged using TissueFAXS fluorescent slide scanner and analyzed using StrataQuest software. Colors denote different cellular markers: blue for Ki67^+^ active proliferating cells, grey for CD35^+^ follicular dendritic cells, yellow for CD3^+^ T cells, and red for IgD^+^ B cell follicles. Scale bar is 500 µm. Error bars display mean ± SD. NS, no significance; *, P<0.05; **, P<0.01; ***, P<0.001; ****, P<0.0001.

To investigate the mechanisms underlying the enhanced antibody response induced by MB-P72, we assessed the follicular T helper (Tfh) cell response in the spleen at day 35. Tfh cells were identified as CD4^+^B220^-^CD44^+^PD-1^+^CXCR5^+^ (Extended Figure 4). The frequencies of Tfh cells were significantly higher in the spleen of MB-P72 immunized mice (6.6%) than S-P72 immunized mice (4.4%), IC-P72* immunized mice (4.8%) and placebo mice (1.6%) (Figure 3F). Spleen sections were stained for IgD to identify B cell follicles, CD35 for follicular dendritic cells, CD3ε for T cells, and Ki-67 for actively proliferating cells. MB-P72 immunization induced significantly more germinal centers (21 per spleen section) compared to immunization with IC-P72* (13 per spleen section) and S-P72 (15 per spleen section) (Figure 3G, H). These findings suggest that the enhanced antibody response observed with MB-P72 immunization is associated with increased germinal center formation and a stronger Tfh cell response.

**Figure 4.**
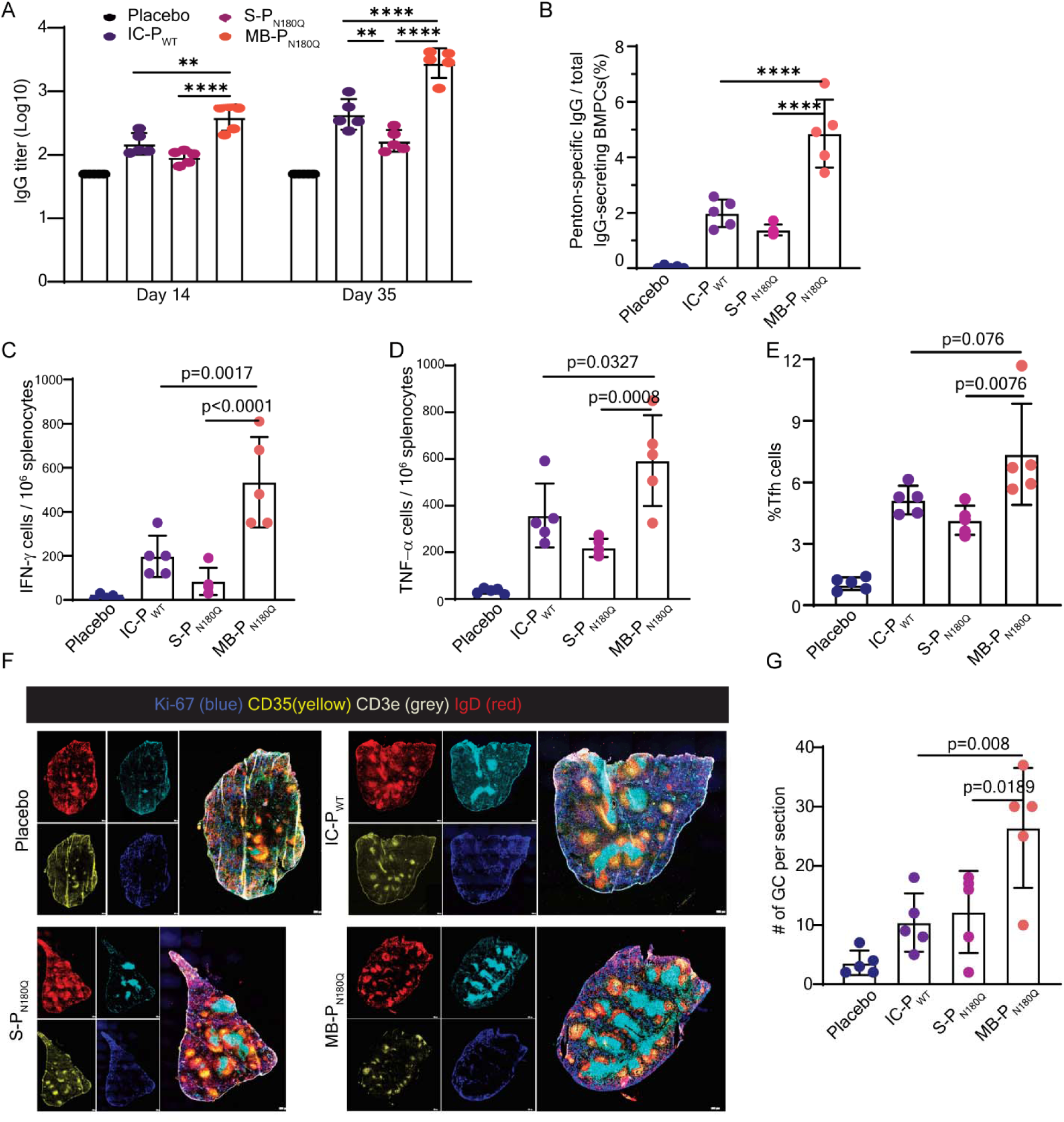
MB-Penton is more immunogenic than S-Penton and IC-Penton in mice. (**A**) Comparison of penton-specific IgG titers at day 14 and day 35 among four groups of mice: placebo (blue), IC-P_wt_ (purple), S-P_N180Q_ (magenta), and MB-P_N180Q_ (orange) (n=5 mice per group). (**B**) The frequencies of penton-specific IgG secreting cells among total IgG-secreting bone marrow cells by ELISPOT (n=5). Plasma cells were enriched from bone marrow using anti-CD138 magnetic beads. 10^4^ enriched cells were incubated with penton coated wells to assay for penton-specific IgG-secreting cells and 10^3^ enriched cells were incubated in wells coated with anti-IgG to assay for total IgG-secreting cells. Data are shown as mean ± SD. (**C, D**) The number of IFN-γ (C) and TNF-α (D) secreting cells (SFC) per 10^6^ splenocytes from different groups (n=5). Splenocytes were stimulated with recombinant penton protein for 36 hours and cytokine-secreting cells were quantified by an IFN-γ/TNF-α dual color ELISpot assay. (**E**) Comparison of percentages of CD4^+^B220^-^CD44^+^PD-1^+^CXCR5^+^ Tfh cells in the spleen of different groups of mice (n=5 per group). Tfh cells were identified the same way as in Figure 3F. (**F**) Representative immunofluorescent images of spleen sections of immunized mice (F) and quantification of the numbers of germinal centers (G) (n=5 mice per group). Germinal centers were stained in the same way as in Figure 3G. One-way ANOVA was used for statistical analysis between the indicated groups. NS, no significance; *, P<0.05; **, P<0.01; ***, P<0.001; ****, P<0.0001.

### MB-Penton is more immunogenic than S-Penton and IC-Penton in mice

We also compared the immunogenicity of IC-, S- and MB-Penton by immunizing BALB/c mice with 5 μg of IC-P_WT_, S-P_N180Q_, and MB-P_N180Q_ mRNA in LNP formulation (using ARV L002 ionizable lipid) at day 0 and 21. Sera was collected before immunization and two weeks after each immunization (days 14 and 35) and used to assay penton-specific IgG by ELISA. The IgG titers were 159 for IC-P_WT_, 95 for S-P_N180Q_, and 427 for MB-P_N180Q_ 14 days after the first immunization (Figure 4A). 14 days after the second immunization, the titers rose to 494 for IC-P_WT_, 177 for S-P_N180Q_, and 3056 for MB-P_N180Q_. Consistently, the frequency of penton-specific ASCs in the bone marrow at day 35 was significantly higher in MB-P_N180Q_ immunized mice than in IC-P_WT_ and S-P_N180Q_ immunized mice (Figure 4B). Similarly, the frequencies of IFN-γ- and TNF-α-secreting T cells following stimulation of splenocytes with recombinant penton protein were significantly higher in MB-P_N180Q_ immunized mice than in IC-P_WT_ and S-P_N180Q_ immunized mice (Figure 3C, D). In a separate experiment where penton mRNAs were formulated in LNP using ionizable lipid SM-102, MB-P_N180Q_ also stimulated higher antigen-specific IgG responses than IC-P_WT_ and S-P_N180Q_ (Extended Figure 3C-E). These results show that MB-P_N180Q_ is more immunogenic than IC-P_WT_ and S-P_N180Q_ in inducing both antibody and T cell responses in mice.

We also investigated the mechanisms underlying the enhanced immune responses induced by MB-P_N180Q_. The frequencies of CD4^+^B220^-^CD44^+^PD-1^+^CXCR5^+^ Tfh cells were significantly higher in the spleens of MB-P_N180Q_ immunized mice (7.4%) than IC-P_WT_ immunized mice (5.1%), S-P_N180Q_ immunized mice (4.1%) and placebo mice (1%) (Figure 4E and Extended Figure 4). Germinal center analysis revealed that MB-P_N180Q_ induced significantly more germinal center formations than IC-P_WT_ and S-P_N180Q_ (Figure 4F). Mice immunized with MB-P_N180Q_ had an average of 26 germinal centers per section, which was 2.2 times higher than in S-P_N180Q_-immunized mice (12 per section, p=0.019) and 2.5 times higher than in IC-P_WT_-immunized mice (10 per section, p=0.008) (Figure 4G). These results show that the enhanced antibody response observed with MB-P_N180Q_ immunization is associated with increased germinal center formation and a stronger Tfh cell response.

### Membrane-bound P72 and penton induce robust antibody and T cell responses in pigs

We further assessed the immunogenicity of the MB-P72 and MB-P_N180Q_ mRNA vaccines in pigs. Five-week-old piglets were injected intramuscularly with 30 μg of mRNA in LNP formulation at day 0 and 21 (Figure 5A). Sera were collected before immunization and weekly following immunization for assaying antigen-specific IgG titers. Spleen were harvested at day 35 for assaying T cell responses. Antigen-specific IgG responses became detectable 14 days after the first immunization, increased steadily afterwards, and reached the highest level at day 35, i.e., 14 days after boost (Figure 5B). Specifically, P72-specific IgG titers were 174 at day 14 and increased to 81359 at day 35; and penton-specific IgG titers were 332 at day 14 and increased to 8086 at day 35. Antigen-specific T cell responses were assessed by ELISPOT following stimulation of splenocytes with a P72 peptide pool or recombinant penton protein. The frequencies of IFN-γ-secreting cells were 438 and 400 per 10^6^ splenocytes from MB-P72 and MB-P_N180Q_ immunized pigs, respectively (Figure 5C). These results show that both MB-P72 and MB-P_N180Q_ induce robust antibody and T responses in pigs.

**Figure 5.**
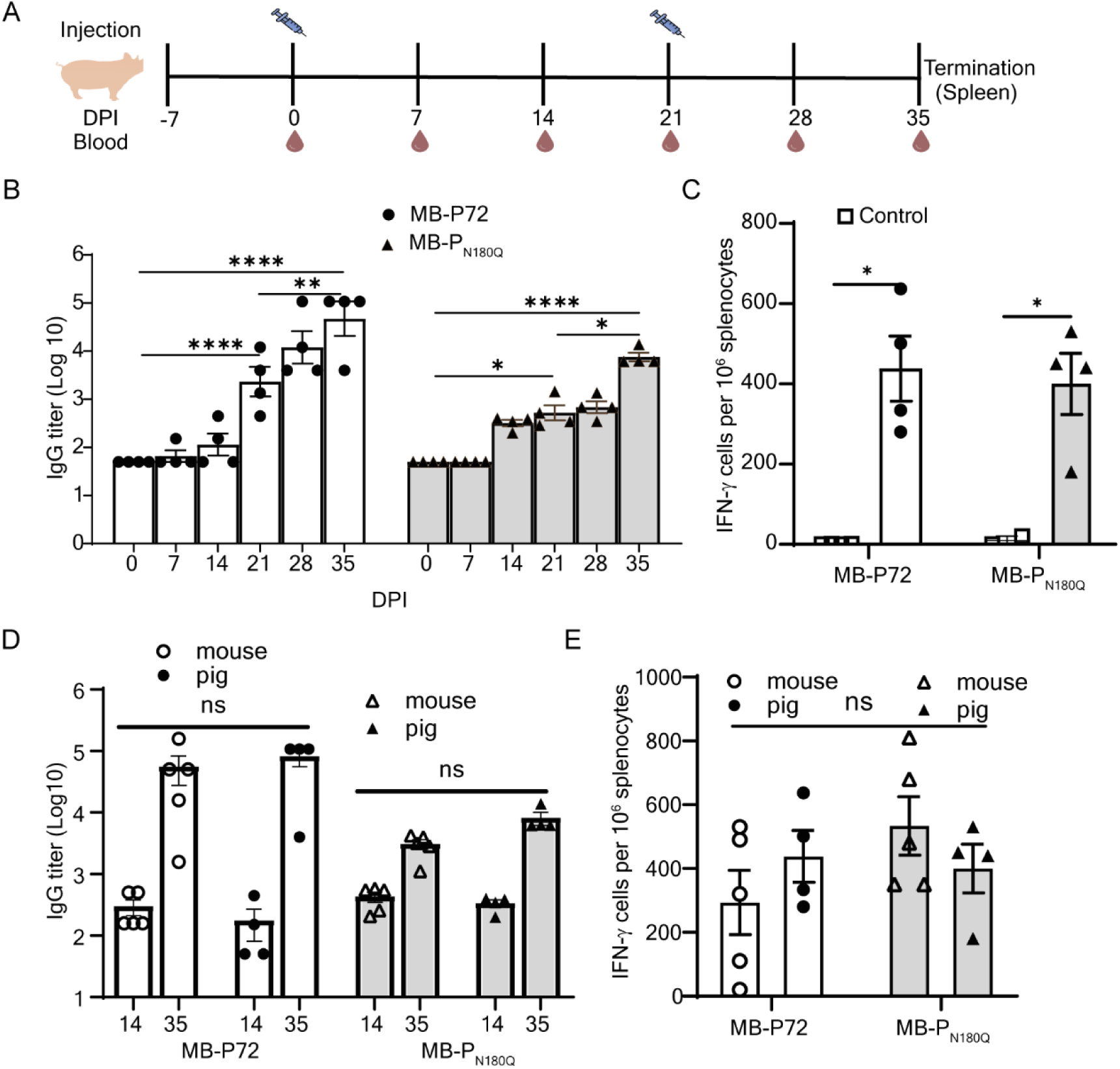
Membrane-bound P72 and penton induce robust antibody and T cell responses in pigs. (**A**) Two groups of pigs (n=4 per group) were immunized with LNP-mRNAs expressing MB-P72 or MB-P_N180Q_, and one group of pigs (n=2) were injected with sterile PBS at day 0 and 21. Sera were collected before immunization and weekly after immunization for assaying antigen-specific IgG responses by ELISA. Spleen was harvested at day 35 and single cell suspension were stimulated with a P72 peptide pool or recombinant penton protein to assay IFN-γ secreting cells by ELISPOT. (**B**) Antigen-specific IgG titers in MB-p72 or MB-P_N180Q_ immunized pigs over time. (**C**) The frequencies of IFN-γ secreting cells (SFC) per 10^6^ splenocytes from MB-P72 or MB-P_N180Q_ immunized pigs. (**D**) Comparison of antigen-specific IgG titers among MB-P72 or MB-P_N180Q_ immunized mice and pigs. (**E**) Comparison of IFN-γ secreting cells among MB-P72 or MB-P_N180Q_ immunized mice and pigs.

We conducted a comparative analysis of MB-P72- and MB-P_N180Q_-induced antibody and T cell responses between mice and pigs. The antigen-specific IgG responses were nearly identical between the two species for both MB-P72 and MB-P_N180Q_ (Figure 5D). For MB-P72-induced T cell responses, IFN-γ levels were lower in mice compared to pigs, whereas for MB-P_N180Q_-induced T cell responses, IFN-γ levels were higher in mice than in pigs (Figure 5E). However, none of these differences were statistically significant. These findings indicate that MB-P72 and MB-P_N180Q_ elicit similar antibody and T cell responses in both mice and pigs, cross-validating the experimental outcomes.

## Discussion

In this study, we combined mRNA vaccine technology with protein engineering to tackle the longstanding challenge of low immunogenicity associated with intracellular viral proteins. Using ASFV capsid proteins P72 and penton as models, we show that when these proteins are expressed as membrane-bound and secreted forms, they fold and assemble into native multimeric structures without the help of viral chaperone. Importantly, both the membrane-bound P72 and penton induced significantly stronger humoral and cellular immune responses in mice than their secreted and intracellular counterparts. Our study provides a simple approach to enhance antigen presentation and immunogenicity of intracellular proteins, opening the possibility for developing safe and effective subunit vaccines.

Our key innovation is to engineer ASFV capsid proteins P72 and penton as membrane-bound and secreted proteins, which facilitated their folding and assembly into native multimeric structures. Naturally, viral capsid proteins are synthesized in the cytosol of infected host cells. With the help of chaperones, the newly synthesized capsid proteins are folded and assembled into intermediate multimeric structures and finally into much larger and more complex capsid, into which viral genome is packaged. The intermediate for P72 is a homotrimer and for penton a homopentamer^11,12^. Studies have shown that folding and trimerization of P72 requires an ASFV-encoded chaperone pB602L^20,21,25^. Consistently, we found that expression, folding and trimerization of the native intracellular P72 requires chaperone pB602L. In contrast, the membrane-bound and secreted P72 were readily detected without pB602L. Furthermore, ∼50% of the membrane-bound and secreted P72 form trimers based on non-reducing gel electrophoresis followed by Western blotting, size exclusion chromatography and transmission electron microscopy of purified P72. As newly synthesized membrane-bound and secreted proteins are directed into the ER and Golgi apparatus, the oxidative environment of the ER likely facilitated the proper folding, disulfide bond formation and trimerization of P72 without the chaperone pB602L^27–31^. In comparison, approximately 30% of both intracellular and secreted penton formed pentamer, but almost 100% of the membrane-bound penton was detected as pentamer. It is possible that the long and flexible hinge used to present penton (and P72) on the cell surface may help to stabilize the multimeric complexes. It is notable that the membrane-bound and secreted penton became glycosylated likely during their transition through the Golgi apparatus. We were able to introduce a single amino acid mutation of N180Q to abolish the glycosylation and therefore to preserve immunogenic potential without masking antigenic epitopes by glycans^32^. The ability to express P72 and penton in their native multimeric structures through a simple engineering of them as the membrane-bound or secreted proteins should help to preserve their conformational epitopes, providing a structural basis for inducing the appropriate antibody responses.

We show that the membrane-bound P72 and penton induced significantly stronger immune responses than their secreted and intracellular counterparts in mice when delivered through mRNA in LNP formulation. Antigen-specific IgG titer induced by MB-P72 was 540-fold higher than that induced by IC-P72* (50,820 vs. 93) 14 days after boost. Consistently, the frequencies of antibody secreting cells in the bone marrow and the frequencies of T cells that were stimulated by P72 peptides to secrete IFN-γ and TNF-α tended to be higher although not statistically significant. MB-P72 also induced potent IgG responses in pigs, reaching a tier over 80,000 14 days after second immunization. Similarly, antigen-specific IgG titer induced by MB-P_N180Q_ was 19-fold higher than that induced by IC-P_WT_ (3056 vs. 494) 14 days after boost. Consistently, the frequencies of antibody secreting cells in the bone marrow and frequencies of T cells that were stimulated by recombinant penton to secrete IFN-γ and TNF-α were all significantly higher following immunization with MB-P_N180Q_ than IC-P_WT_. MB-P_N180Q_ also induced potent IgG responses in pigs, reaching a titer over 8,000 14 days after second immunization. The enhanced antibody responses are associated with elevated Tfh response and germinal center formation in the spleen. Our results are consistent with previous observation showing that the membrane-bound Spike protein induced stronger antibody response than the secreted form of Spike protein^33,34^. Multiple factors may have contributed to the enhanced immunogenicity of the membrane-bound P72 and penton. First, a large number of the membrane-bound P72 and penton are displayed on the cell surface. The increased epitope density and their readily accessible to B cell recognition likely help to stimulate antibody responses. Second, the formation of trimeric P72 and pentameric penton when expressed in the membrane-bound form preserves conformational epitopes, which could stimulate additional antibody responses. Third, a higher level of the membrane-bound P72 or penton may be expressed following LNP mRNA vaccination. The induction of significantly stronger antibody and T cell responses by both the membrane-bound P72 and penton, which differ in size and multimeric configuration, suggests our finding could be a generally phenomenon.

The success of our approach with ASFV capsid proteins highlights its potential for developing safe and effective subunit vaccine for ASFV as well as application to other intracellular viral antigens. Many viral capsid proteins face similar challenges in antigen accessibility and presentation. By applying protein engineering to enhance extracellular presentation on the membrane, coupled with the flexibility of mRNA vaccination, our simple approach could be a powerful tool for developing subunit vaccines against a broad spectrum of intracellular pathogens, without resorting the use of artificial multimeric structures such as foldon^35^. Our approach enables the targeting of intracellular viral proteins that have historically been difficult to use as immunogens due to their reliance on specific intracellular pathways and post-translational modifications for folding and multimeric formation.

Our study has two specific limitations: First, we did not assay neutralizing activity of antibodies induced by the membrane-bound, secreted and intracellular forms of P72 and penton, therefore we do not know if antibodies induced by the membrane-bound P72 and penton are more effective in neutralizing ASFV due to responses to conformational epitopes. This is because antibodies elicited by a single ASFV antigen are known not to give reliable neutralization results due to multiple entry pathways used by infectious ASFV virions^36–41^. Both the extracellular enveloped virion, i.e., with the outer membrane, and intracellular mature virion, i.e., without the outer membrane, are infectious^42^. Furthermore, ASFV can infect host macrophages through multiple entry mechanisms, including receptor-mediated endocytosis^36,37^, clathrin-mediated endocytosis^38,40^, macropiniocytosis^39,41^, and phagocytosis^41^. Second, we did not compare the immunogenicity of the three different forms of P72 and penton in pigs due to cost consideration. Nevertheless, our study of combining mRNA vaccine technology with protein engineering provides a simple approach for enhancing folding and formation of native multimeric structures of capsid proteins for ASFV subunit vaccine development as well as enhancing immunogenicity of intracellular proteins in general.

## Methods and Materials

### Animal Experiments

All procedures were conducted in accordance with the approved animal protocol 0322-021-25, which had been reviewed and approved by the Massachusetts Institute of Technology (MIT) Committee on Animal Care. The experiments adhered to the established guidelines for animal care. Female BALB/c mice, 7 weeks old, were procured from Charles River Laboratories and were housed within an MIT animal facility. LNP-mRNA vaccination of pigs was performed according to the protocols approved by Committee on Animal care (protocol number 2308000566) and Midwest Veterinary Service (MVS), Inc. (protocol number 24005).

### Cells and Plasmids

HEK293T (ATCC CRL-3216) cells were purchased from the American Type Culture Collection (ATCC) and grown in DMEM supplemented with 2 mM L-glutamine, non-essential amino acids, 100 U/mL gentamicin, and 10 % fetal bovine serum (FBS) (Invitrogen Life Technologies). Vero (ATCC CCL-81) and MCDK (ATCC CCL-34) cells were purchased from the American Type Culture Collection (ATCC) and grown in Eagle’s Minimum Essential Medium (EMEM), 10 % fetal bovine serum (FBS) (Invitrogen Life Technologies) and 1 X pen/strep. 3D4/31 (ATCC CRL-2844) cells were purchased from ATCC and grown in RPMI-1640, 10 % filter sterilized FBS (ATCC), and 1 X pen/strep. Gene information of ASFV viral protein P72, penton, P17, and pB602L was obtained from ASFV Georgia 2007/1 (Gene bank # FR682468.2). ASFV DNA encoding for three forms (membrane-bound, secreted, and intracellular) of P72 and penton, pB602L, and P17 were codon optimized to swine and synthesized by GenScript (Piscataway NJ) and cloned into phCMV1 and/or phCMV3 (HA-tagged) (Promega) via the NEBuilder® HiFi DNA Assembly (NEB, #M5520AVIAL), and the corresponding ASFV plasmids were constructed for the following characterization. Secreted form of capsid proteins was engineered by addition of signal peptide from a human CD8α (GenBank ID: NP_001139345.1) to the N terminus, while the membrane-bound form encompasses an additional CD8α stalk region or hinge, transmembrane region, and short cytoplasmic tail to the C terminus.

### Structural Modeling

The three-dimensional (3D) structures of both capsid protein subunit and multimeric forms (trimer for P72, pentamer for penton) were predicted by AlphaFold2^23^, followed by structural visualization using the open-source PyMOL system (PyMOL Molecular Graphics System, version 1.7, Schrödinger, LLC). Predicted structures of engineered antigens with top-ranked models were aligned with the intracellular or native form of each capsid protein for similarity analysis by calculation of the root-mean-square deviation (RMSD) metric score (the lower the score the more similar the structures).

### Transfection of Plasmid Expressing ASFV Viral Protein in Mammalian Cells

Confluent monolayers of HEK293T, Vero, and 3D4/31 cells were transfected with constructed plasmids using Lipofectamine 3000 transfection reagent (Invitrogen, #L3000008) according to manufacturer’s protocol. Forty-eight hours after transfection, cells or the whole cell lysate were used for further analysis.

### Confocal microscopy

#### Intracellular staining

Preconfluent monolayers were rinsed with phosphate-buffered saline (PBS), fixed with 4% paraformaldehyde at room temperature for 20 min, washed three times with PBS and rinsed for 5 min after each wash, permeabilized with 0.1% Triton X-100 in phosphate-buffered saline (PBS-T) at 4 °C for 10 min, washed with PBS for 5 min at room temperature, blocked with PBS-B solution (PBS containing 4% BSA) at 37 °C for 30 min and incubated with mouse anti-P72 antibody (MyBioSource’s), anti-HA antibody, or anti-His antibody at room temperature for 1 h, followed by three washes with PBS. Then, the cells were incubated with fluorescein conjugated (AF488, CF568) goat anti-mouse antibody at room temperature in dark for 1 h, followed by 1 μg/mL 4 ′, 6-diamidino-2-phenylindole dihydrochloride (DAPI) solution incubation for 15 min. Cells were then washed three times as described above and observed using a fluorescence microscope (Olympus FV1200 Laser Scanning Confocal Microscope).

#### Surface Staining

Preconfluent monolayers were rinsed with phosphate-buffered saline (PBS), blocked with PBS-B solution (PBS containing 4% BSA) at 37 °C for 30 min and incubated with mouse anti-P72 antibody (MyBioSource’s), anti-HA antibody, or anti-His antibody on ice for 1 h, followed by three washes with PBS. Then, the cells were incubated with fluorescein conjugated (CF568) goat anti-mouse antibody on ice in dark for 1 h, followed by 1x CellMask™ Green (Invitrogen, # C37608) and 1 μg/mL DAPI solution incubation for 15 min. Cells were then washed three times as described above and observed using a fluorescence microscope (Olympus FV1200 Laser Scanning Confocal Microscope).

### SDS-PAGE and Western Blotting Analysis

The transfected cells were harvested and lysed using RIPA lysis buffer (Thermo Fisher, #89900) following the manufacturer’s instructions. Subsequently, the lysed samples were combined with one-third of the total volume of 4 × sample loading buffer, with or without SDS or β-mercaptoethanol, and boiled for 10 min. To detect native proteins under non-reducing conditions, cleared cell lysate was incubated with 4× NuPAGE LDS sample buffer (Thermo Fisher Scientific) without heat-denaturing and β-mercaptoethanol reducing. To detect N-linked glycosylation of penton, transfected cell lysates were denatured first, then incubated with PNGase F (New England Biolabs) at 37 °C for 1 h. The protein samples were resolved by SDS-PAGE using gels of varying percentages (8%, 10%, or 12%). After electrophoresis, the separated proteins were transferred onto a nitrocellulose membrane, which was subsequently blocked using 5% skimmed milk at 4 °C overnight. Next, the membranes were subjected to immunoblotting using specific primary antibodies, including anti-P72, anti-HA, anti-His, and anti-actin or anti-tubulin antibodies, at room temperature for 1 h. The membranes were washed three times with PBST (Phosphate Buffered Saline with 0.05% Tween-20) and then incubated with anti-mouse HRP-conjugated secondary antibodies (Thermo Fisher) for 1 h at room temperature. Subsequently, the membranes were washed three times with PBST. The protein bands were visualized using the Bio-Rad system (Bio-Rad, USA), and the intensities of the bands were quantified using Image Lab imaging software (Bio-Rad).

### Flow Cytometry Analysis

Transfected cells expressing ASFV viral proteins were detached using trypsin, mixed with an excess of PBS, and centrifuged at 1200 rpm for 10 min. The cells were washed once with FACS buffer (PBS, 2% BSA) and resuspended in 200 µL of FACS buffer. To minimize nonspecific binding of mouse monoclonal antibodies, the Rhesus Fc Receptor Binding Inhibitor was added to the samples. The samples were then incubated on ice for 20 min without additional washing. Subsequently, without washing the samples, primary antibody staining was performed using a 1:1000 dilution of mouse anti-P72 or anti-HA antibody for 30 min on ice. After staining, the cells were washed three times with FACS buffer. Next, a 1:500 dilution of anti-mouse-AF647 secondary antibody was applied, and the cells were incubated for 30 min on ice, followed by three washes with FACS buffer. Finally, just before analysis, 5-10 µL of propidium iodide (PI) staining solution was added to each sample. The stop count for analysis was set based on viable cells as determined from a dot-plot of forward scatter versus PI.

### Recombinant Protein Production and Sample Preparation for P72 Trimer Characterization

The synthesized genes were cloned into the phCMV3 vector separately. A C-terminal HA tag was included in this vector. HEK293T cells were cultured with DMEM plus 10 % FBS at 37 °C and 5 % CO_2_. For a 60 mm cell culture dish, the cells were transfected with plasmids containing 10 μg MB-P72 or S-P72, 10 μg IC-P72 and 5 μg pB602L plasmid mixture and 30 or 45 μg PEI well mixed in Opti-MEM (Gibco, #31985062). The transfected cells were harvested 48 h post-transfection with RIPA buffer and cocktail protease inhibitors (Thermo scientific, # A32965) for 30 min on ice then scraped off the dish. The whole cell lysate was sonicated for 2 min and the cell lysate was centrifuged for 60 min at 20,000 rpm (JA 25.50 rotor, Beckman). The recombinant P72 protein in the supernatant were collected and applied to the anti-HA Magnetic Beads (MCE, #HY-K0201). The bound protein was washed with the resuspended buffer five times and then was eluted from the beads with a buffer containing 2 mg/mL HA peptide. The eluted sample was concentrated and further purified by Superdex 200 increase 10/300 GL size exclusion column (Cytiva 28990944) via capillary loop running in a buffer containing 20 mM HEPES at pH 7.4 and 300 mM NaCl. The P72 protein from size exclusion column peak was concentrated and kept at −80 °C for future use.

In sample preparation for negative stained-electron microscopy (TEM), 10 µL of protein sample and buffer containing solution was dropped on a 200 meshes copper grid coated with a continuous carbon film and waited for 60 sec and removed excess solution by touching the grid with a kimwipes and then 10 µL of negative staining solution, phosphotungstic acid, 1% aqueous solution was dropped on the TEM grid and immediately removed it by kimwipes and 10 µL of the stain was then applied to the grid and after 30 sec, the excess stain was removed by touching the edge with kimwipes. Finally, dried the grid at room temperature. After that, the grid was mounted on a JEOL single tilt holder equipped in the TEM column. The specimen was cooled down by liquid-nitrogen and imaging on a JEOL 2100 FEG microscope was done using minimum dose method that were essential to avoid sample damage under the electron beam. The microscope was operated at 200 kV and with a magnification in the ranges of 10,000∼60,000 for assessing particle size and distribution. All images were recorded on a Gatan 2k x 2k UltraScan CCD camera.

### Production of mRNA and LNP Formulation

Genes encoding IC-P72, S-P72, MB-P72, pB602L, IC-P_WT_, S-P_N180Q_ and MB-P_N180Q_ were inserted into pUC57 (primers are listed in Extended Table 1), which contains a T7 promoter, 5UTR, 3’UTR, and polyA tail. The linearized plasmids were subject to in vitro transcription using pseudo-UTP. After mRNA purification and capping reaction, an aqueous phase of mRNA was prepared by diluting mRNA stock in 10 mM citrate buffer. The organic phase of lipid nanoparticles was prepared by adding 200 proof ethanol with lipid stock solutions which contained ionizable lipid SM102 or L002 (Advanced RNA Vaccine Technologies, ARV), DSPC helper lipid (Avanti Polar Lipids), cholesterol (Avanti Polar Lipids), and DMG-PEG-2000 (Avanti Polar Lipids). LNP-mRNA formulations were prepared by mixing organic and aqueous phases at a ratio of 1:3. RiboGreen assay (Thermo Fisher) was performed by following manufacture’s instruction to quantify mRNA after formulation. Encapsulation efficiency was measured by Picogreen assay. To validate LNP delivery of mRNAs, HEK293T and Vero cells were seeded onto 24-well tissue culture plate and 500 ng of LNP-mRNA diluted in Opti-MEM was added in individual well. Cells were collected in 48 h and protein expression was analyzed by flow cytometry and Western blotting as described below.

### Expression of ASFV mRNA *in vitro*

ASFV mRNAs were transfected into HEK293T and Vero cells with MessengerMax (Invitrogen) for naked mRNA or formulated mRNA-LNP directly. After 48h, the expression was evaluated with flow cytometry (FACS), immunofluorescence assay (IFA), and western blot (WB). For IFA, cells were incubated with mouse anti-P72 antibody or mouse anti-HA antibody and AF488 conjugated anti-mouse secondary antibody (Abcam), and images were taken with a fluorescence microscope (Olympus FV1200 Laser Scanning Confocal Microscope). For FACS, cells were trypsinized and incubated with mouse anti-P72 antibody or mouse anti-HA antibody and AF488 conjugated anti-mouse secondary antibody (Abcam). Data was acquired with LSR Fortessa HTS-2 (BD Biosciences) and analyzed with FlowJo (BD Biosciences). For WB, cells were harvested and denatured in lysis buffer. Samples were loaded and run in 8 or 12 % SDS-PAGE gel and transferred to nitrocellulose membrane. The membrane was incubated with mouse anti-P72 antibody or mouse anti-HA antibody and HRP conjugated anti-mouse secondary antibody (Invitrogen, Cat#62-6520). Anti α-actin HRP Antibody for protein loading control was purchased from Santa Cruz Biotechnology (sc-47778 HRP).

### Immunization of Mice

To assess immunogenicity of individual antigens in mice, six to eight-week-old female BALB/c mice were purchased from Charles River and housed in animal facility at MIT. Briefly, 5 mice were assigned to each group and immunized intramuscularly with 50 µL 5 µg LNP-mRNA (SM102 or ARV-L002) diluted in PBS. All mice were boosted three weeks later. Serum samples were collected from submandibular vein prior to immunization and two weeks after each injection. Mice were euthanized at day 35 and spleen tissue and bone marrow were collected for analysis of cellular immunity.

### Immunization of Pigs

A total of ten four-week-old piglets were randomly assigned to three groups: four piglets each in the MB-P72 and MB-P_N180Q_ groups, and two piglets in the control group, which received sterile PBS. 30 μg of LNP-mRNA expressing each individual antigen was diluted to 1 mL in sterile PBS and injected to the back of ear intramuscularly and boosted three weeks later. Serum samples were collected before vaccination and weekly after each injection. Body weight of each pig was measured before the study and at the termination. All pigs were euthanized two weeks after boost and spleens were collected for testing T cell responses.

### Antibody Measurement with ELISA

Each well of a flat-bottomed, high-binding 96-well plate (Santa Cruz Biotechnology) was coated with approximately 50 μL of antigen solution, prepared at a concentration of 5 μg/mL of purified recombinant ASFV viral protein in PBS (pH 7.4). The plates were then incubated at 4 °C overnight. Following the incubation, the antigen solutions were carefully aspirated, and the plates underwent five washes using PBS Tween (PBST, containing 0.05% v/v Tween-20; pH 7.4). Subsequently, the wells were filled with blocking buffer (5% skimmed milk in PBS) and incubated at 37 °C. After a 1 h incubation at room temperature, the blocking buffer was aspirated, and all wells received an additional five washes with PBST. For the subsequent step, mouse sera were subjected to three-fold serial dilutions, starting at a 1:100 dilution, in PBS containing 5% skimmed milk. These diluted sera were added to the plates at a volume of 100 μL per well and incubated for 1 h at 37 °C. Following this incubation, the plates were washed five times with PBST. To detect IgG, goat anti-mouse IgG conjugated to HRP (Cell Signaling Technology, diluted 1:2,000 in PBS with 5% skimmed milk) was used as the secondary antibody. The secondary antibody solution was added to the plates, and the plates were incubated at 37 °C for 1 h. Afterward, the plates underwent five additional washes using PBST. To develop the assay, 100 μL of HRP substrate 3,3’,5,5’-tetramethylbenzidine (TMB) (Cell Signaling Technology) was added to each well. The plates were gently shaken for 15 min, and the reaction was stopped by adding 100 μL of a stop solution (0.2 M H_2_SO_4_) to each well. The absorbance was measured at two wavelengths, 450 nm (signal) and 570 nm (background), using a Tecan microplate reader. End-point titers were determined as the highest serum dilution at which the optical density difference, compared to serum from sham-vaccinated mice at the same dilution, exceeded 0.005.

### Enzyme-linked Immunospot (ELISPOT) assay

Splenocytes from vaccinated mice were evaluated for antigen specific IFN-γ by Enzyme-linked immunospot (ELISPOT). ELISPOT assays were performed using a Mouse IFN-γ or TNF-α Single-Color ELISPOT kit from ImmunoSpot followed the manufacturer’s instruction. Briefly, splenocytes were plated at 4 x 10^5^ cells/well and co-cultured with either 5 μg/mL P72 peptide pool (Core facility, Koch Institute), 2 μg/ mL recombinant penton protein, Cell Stimulation Cocktail (500X) containing phorbol 12-myristate 13-acetate (PMA) and ionomycin (eBioscience, # 00-4970-93) or medium alone in a total volume of 200 μL/well T cell media for 36 h at 37 °C in 5 % CO_2_. The plates were incubated with detection antibodies, Biotin-IFN-γ and Streptavidin-HRP at RT for 1-2 h, respectively. The plates were developed with 50 μL/well the development solution for up to 30 min. Color development was stopped by washing under running tap water. After air-dried, colored spots were counted using a CTL ELISPOT Reader System and CTL ELISPOT Software.

For B cell ELISpot, bone marrow from mouse femur and tibia of both hind limbs were collected into PBS supplemented with 1 mM EDTA and 2% FBS. Bone marrow plasma cells (BMPCs) were further enriched using EasySep™ Release Mouse CD138 Positive Selection Kit (StemCell Technologies) and cryopreserved in 10% DMSO in FBS. A mouse IgG single-color ELISpot kit (ImmunoSpot) was used to determine total IgG-secreting cells in enriched BMPCs according to the manufacturer’s instructions. ASFV proteins (300 ng/well) were coated onto the ELISpot plate to determine the antigen-specific-binding IgG-secreting cells in BMPCs. ELISpot plates were further analyzed using ELISpot plate reader (Cellular Technology).

### Germinal Center Reactions

To image the germinal centers, one fifth of the spleen was embedded in Optimal Cutting Temperature (O.C.T.) solution (VWR) solution for 5 min at room temperature followed by snap-freezing in liquid nitrogen. Tissue blocks were stored in −80 °C until cryostat sectioning into 6 μm sections using a microtome. Sectioned tissue slides were first air dried for 30 min and an unbroken circle was drawn with a PAP pen (Vector Laboratories) to create a hydrophobic barrier. The slides were blocked in room temperature (RT) for 2 h by adding PBS containing 2% BSA, 10% normal goat serum, and TruStain FcX™ (anti-mouse CD16/32) Antibody (Biolegend). Slides were washed three times with 0.5% Tween-20 in PBS and permeabilized using 2% Trion X-100 (Sigma) followed by staining according to a modified procedure published previously^43^. Briefly, slides were stained with a primary antibody cocktail containing AF647-labled rat anti-mouse IgD antibody (clone 11-26c.2a, Biolegend), Hamster anti-Mouse CD3ε (Clone 500A2, BD Biosciences), biotin-conjugated CD21/CD35 antibody (Clone 8D9, Invitrogen), AF488-conjugated Ki-67 antibody (Clone SolA15, Invitrogen). After overnight incubation at 4°C, slides were washed with PBST for three times followed by adding secondary antibody cocktail containing AF594-conjugated goat anti-hamster IgG (Jackson ImmunoResearch) and AF546-conjugated Streptavidin (Invitrogen). Slides were washed three times and treated with a ReadyProbes™ Tissue Autofluorescence Quenching Kit (Invitrogen) and coverslips were mounted using SlowFade™ Diamond Antifade Mountant (Invitrogen). Mounted slides were protected from light and kept at 4 °C until imaging using TissueFAXS SL Fluorescent Slide Scanner with 20x objective and images were analyzed using StrataQuest software (TissueGnostics).

### Follicular T helper Cell Response

Isolated splenocytes were treated with ACK red blood cell lysis buffer (Gibco) and blocked for Fcγ receptor blockade using TruStain FcX™ (anti-mouse CD16/32) Antibody (Biolegend). Cells were stained by an antibody cocktail containing PE Rat Anti-Mouse CD279 (PD-1) (Clone RMP1-30, BD bioscience), FITC anti-mouse CD4 (Clone GK1.5, Biolegend), PE/Cy7 anti-mouse/human CD44 (Clone IM7, BioLegend), BUV395 Rat anti-mouse CXCR5 (Clone 2G8, BD Bioscience). Stained cells were washed three times and stained for viability by DAPI. Samples were analyzed by flow cytometry on a BD Symphony A3 and analyzed on FlowJo software.

### Statistical Analysis

GraphPad Prism 8.0 was used for conducting statistical analyses and generating data plots. Two-tailed Student’s t-tests were used to assess distinctions between two independent sample groups. One-way analysis of variance (ANOVA) was used for comparisons involving multiple sample groups. The results are presented in the form of mean values along with their standard deviations (mean ± SD). A threshold of P < 0.05 was considered statistically significant. Significance levels were defined as follows: ns (not significant) for P > 0.05; **** for P < 0.0001; *** for P < 0.001; ** for P < 0.01; and * for P < 0.05.

## Acknowledgements

The authors would like to thank Koch Institute Swanson Biotechnology Center and core facilities for assistance with Microscopy, Nanotechnology Materials, Flow Cytometry for data acquisition and analysis, Histology for tissue sectioning, and Biopolymers & Proteomics for peptide synthesis. The authors would also like to thank members of Chen Lab for their suggestions.

## Author contributions

Conceptualization: J Cui, FY, J Chen

Methodology: J Cui, FY, JQ, JHJ, DSY, TW

Investigation: J Cui, FY, J Chen

Supervision: J Chen, RX, HC

Writing—original draft: J Cui

Writing—review & editing: J Cui, J Chen

