## Extended data1 for "Membrane Expression Enhances Folding, Multimeric Structure Formation, and Immunogenicity of Viral Capsid Proteins"

Junru Cui *et al.*

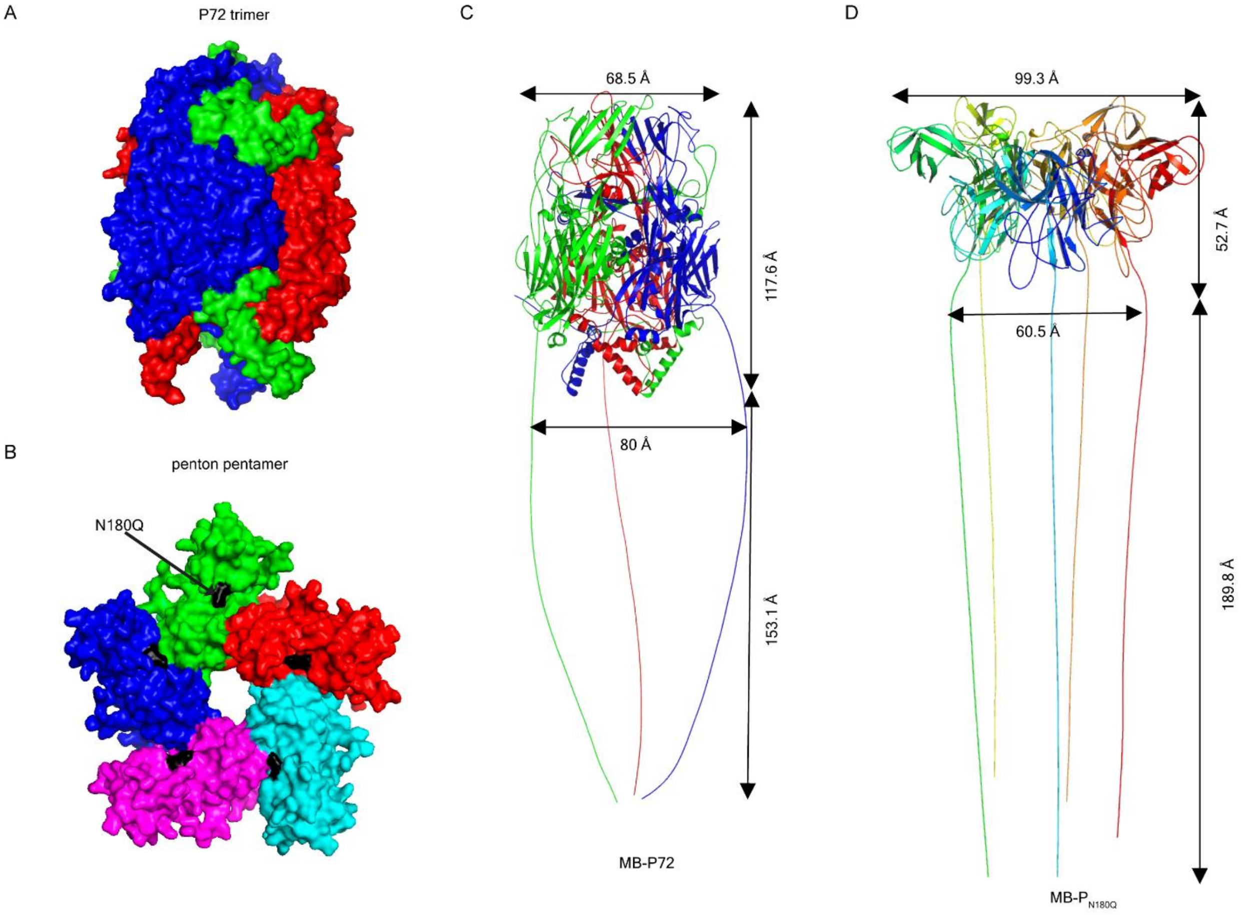


Extended Figure 1. AlphaFold2 modeling of P72 and penton without and with the hinge domain. (**A**) The native trimeric P72 with each monomer in different colors. (**B**) The native pentameric penton with each monomer in different colors. The arrows point to asparagine 180 (N180) that could be glycosylated when expressed as the membrane-bound and secreted forms. (**C**) Three copies of P72-hinge were imported to AlphaFold2 and the structure was visualized using PyMol. (**D**) Five copies of penton-hinge were imported to AlphaFold2 and the structure was visualized using PyMol. The indicated two-dimensional distances (Å) in C and D were measured with measurement wizard function of PyMol.

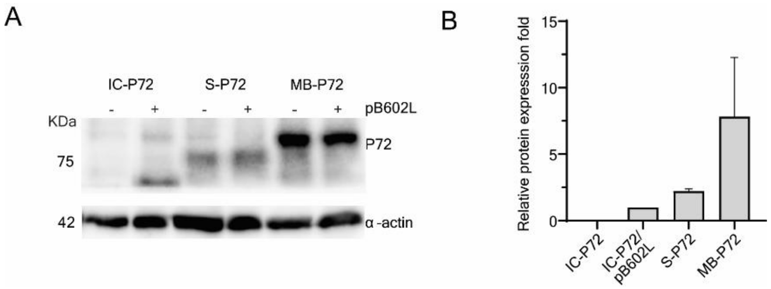


Extended Figure 2. Expression of IC-, S- and MB-P72 from mRNA encoding three forms of P72. (A) Western blotting analysis of P72 protein levels. HEK293T or Vero cells were transfected with mRNA encoding IC-, S- and MB-P72 using MessengerMax$^{TM}$. Cells were lysed 48 hours after transfection and cell lysates were used for Western blotting analysis using mouse anti-P72 followed by HRP-conjugated goat anti-mouse antibodies. (A) A representative WB image. Anti α-actin is used as loading control. (B) Summary of results from three independent experiments shown as mean ± standard deviation (SD) normalized to α-actin levels.

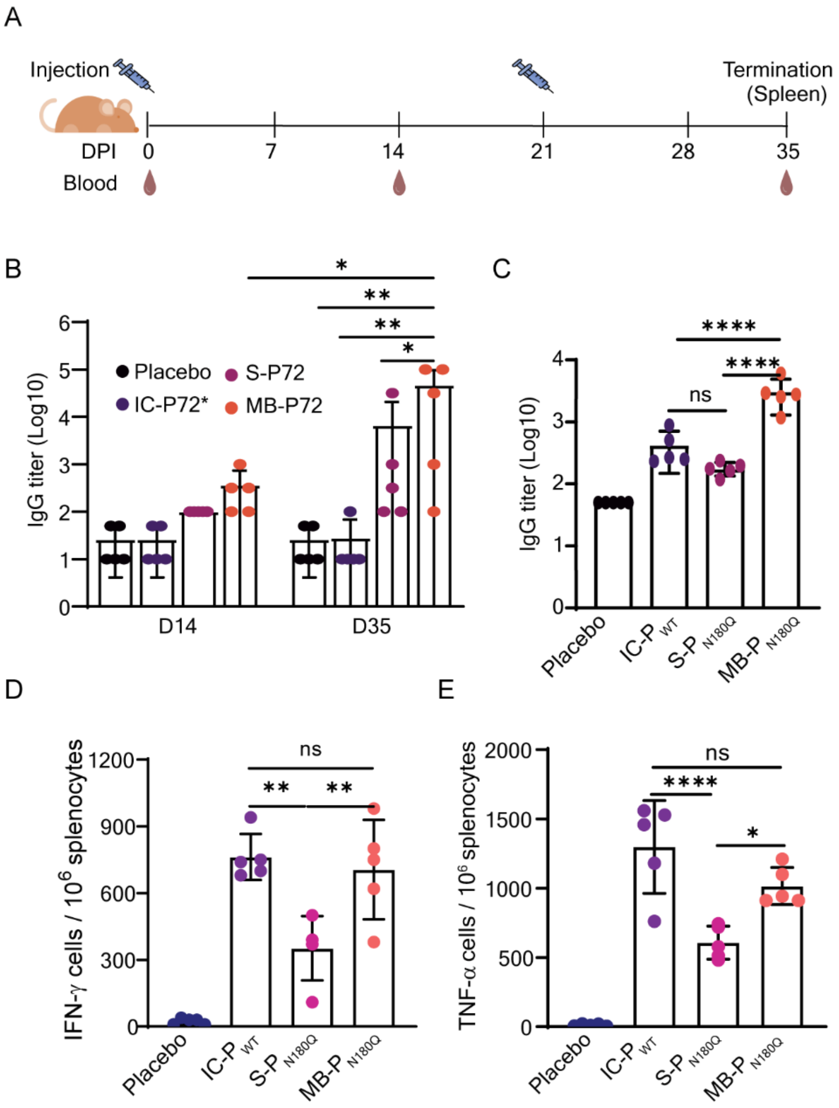


Extended Figure 3. The membrane-bound P72 and penton elicited higher levels of IgG responses than the corresponding IC- and S- counterparts. (**A**) Scheme of immunization. Mice (n=5 per group) were intramuscularly injected at day 0 and 21with mRNA encoding the three forms of P72 and penton formulated in LNP using SM102 ionizable lipid. Sera were collected before priming and 14 days after each immunization and assayed for P72- or penton-specific IgG by ELISA. Splenocytes collected at day 35 were stimulated with recombinant penton protein to assay for IFN-γ and TNF-α-secreting T cells by ELISPOT. (**B**) P72-specific IgG titers on day 14 and day 35. Placebo (blue), IC-P72* (purple), S-P72 (magenta), and MB-P72 (orange) (n=5). (**C**) Penton-specific IgG titers at day 35. Placebo (blue), IC-$P_{WT}$ (purple), S-$P_{N180Q}$ (magenta), and MB- $P_{N180Q}$ (orange) (n=5). (**D, E**) Comparison of the numbers of IFN-γ (D) and TNF-α (E) secreting cells per $10^6$ splenocytes from different forms of penton immunized mice. One-way ANOVA was used for statistical analysis between different groups. NS, no significance; *, P<0.05; **, P<0.01; ***, P<0.001; ****, P<0.0001.

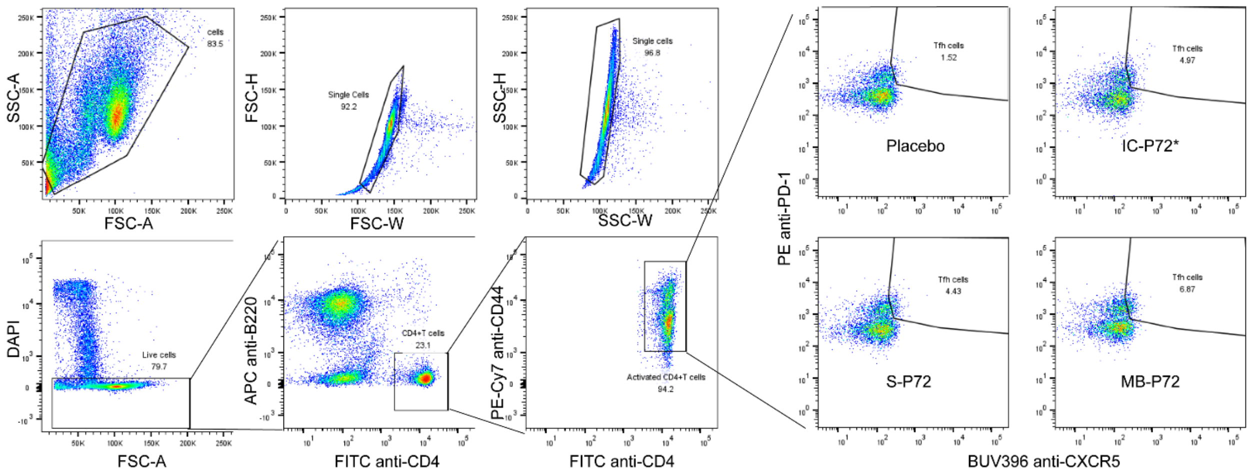


Extended Figure 4. Gating strategy for identifying Tfh cells. Splenocytes from immunized mice were stained with antibodies specific for CD4, B220, CD44, PD-1, and CXCR5 followed by flow cytometry. Single live cells were gated to identify $CD4^{+}B220^{-}CD44^{+}$ activated T cells. Tfh cells were identified as $PD\text{-}1^{+}CXCR5^{+}$ population among the activated T cells.

Extended Table 1. Primers and Peptides used in this study

| |
|---|
| **mRNA constructs** |
| Primers for amplifying various forms of P72 and penton for cloning into pUC57 |
| 1. pUC57 Vector for MB-72 FWD: TCGTCGTTAAGTACGGCTGGAGCCTCGGT |
| 2. pUC57 Vector for MB-/S-P72 REV: AAGGCCATGTGGCTGTTTGAGGTTGCTAGTGAAC |
| 3. Insert MB-/S-P72 FWD CAGCCACATGGCCTTGCCAGTGACAG |
| 4. Insert MB-P72 REV AGCCGTACTTAACGACGATTACGGTGATTGTACAG |
| 5. pUC57 Vector for S/IC-P72 FWD CTCCACCTAAGTACGGCTGGAGCCT |
| 6. pUC57 Vector for IC-P72 REV CTAGCCATGTGGCTGTTTGAGGTTGCTAGTGAACAC |
| 7. Insert S/IC-P72 REV AGCCGTACTTAGGTGGAGTAGCGCAGCAC |
| 8. pUC57-IC-pB602L Vector FWD AACTGTGAGTACGGCTGGAGCCTCGGT |
| 9. pUC57-IC-pB602L Vector REV TCTGCCATGTGGCTGTTTGAGGTTGCTAGTGAACAC |
| 10. Insert IC-pB602L FWD CAGCCACATGGCAGAATTCAATATTGACGAGCTG |
| 11. Insert IC-pB602L REV AGCCGTACTCACAGTTCTGCCTTGGTGTACAG |
| 12. pUC57-MB-$P_{N180Q}$/S-$P_{N180Q}$-F: TTCACTAGCAACCTCAAACAGCCACCATGGGCCACCATGGCCTTGCCAGT |
| 13. pUC57- MB-$P_{N180Q}$-R: AGGCCACCGAGGCTCCAGCCGTACGCTATTAACGACGATTACGGTGATTGT |
| 14. pUC57-IC-$P_{WT}$-F: CACTAGCAACCTCAAACAGCCACCATGGGCCACCATGGCCGCGAATATCATTGC |
| 15. Universal reverse primer for pUC57 HA tag: AGGCCACCGAGGCTCCAGCCGTACGCTATTAAGCGTAATCCGGAACATCG |
| **Eukaryotic expression** |
| 16. IC-P72 FWD: CGCCACCATGGCATCCGGCGGAG |
| 17. IC-P72 REV: GCTCCAGCGGTGGAATACCTTAGTACTGCACTGC |
| 18. S-P72 FWD: TCGACTTAACGGAAATAAGAGAGAAAAGAAGAGTAAGAAGAAATATAAGACCC |
| 19. S-P72 REV: GCTGGTGTGGTAGTAGGCTTGGTGGAATACCTTAGTACTGCACTG |
| 20. pB602L FWD: GCCAAGCTTGCCACCATGGCAGAATTCAATATTGAC G |
| 21. pB602L REV: ATTGGTACCTCACAGTTCTGCCTTGGTGTAC |
| 22. pHCMV3- IC-$P_{WT}$ F EcoRI: ATTGAATTCGCCACCATGGCCGCGAATATCATTGC |
| 23. pHCMV3- IC-$P_{WT}$ /S-$P_{WT}$-HA R BamHI: GCCGGATCCCTTAGAAGTTTTCAGCGAC |
| 24. pHCMV3- S-$P_{WT}$ /MB-$P_{WT}$ F EcoRI: ATTGAATTCGCCACCATGGCCTTGCCAGTGACAG |
| 25. pHCMV3- MB-$P_{WT}$-HA-R: AGCGGATCCACGACGATTACGGTGATTGT |
| **P72 peptide pool**<br>26. SSYIPFHYGGNAIK<br>27. SRISNIKNVNKSY<br>28. SLDEYSSDVTTL |

Extended Table 2. SEC analysis of purified S-P72 and MB-P72

| protein | Peak | Retention (ml) | Area (ml*mAU) | Area % |
|---|---|---|---|---|
| S-P72 | Trimer | 7.867 | 7.769 | 50.77 |
| | Monomer | 21.720 | 5.753 | 37.59 |
| MB-P72 | Trimer | 7.842 | 5.720 | 36.78 |
| | Monomer | 21.703 | 7.489 | 48.15 |

Extended Table 3. Characterization of formulated LNP mRNA

| **Formulation** | **Hydrodynamic size(nm)** | **Polydispersity index** | **Encapsulation efficiency (%)** |
|---|---|---|---|
| IC-P72+pB602L | 112.57 ± 2.056 | 0.123 ± 0.0158 | 91.25 ± 0.026 |
| S-P72 | 114.09 ± 2.533 | 0.090 ± 0.0206 | 93.59 ± 2.646 |
| MB-P72 | 115.33 ± 1.578 | 0.102 ± 0.0085 | 93.88 ± 0.527 |
| IC-$P_{WT}$ | 120.36 ± 3.485 | 0.078 ± 0.0121 | 95.15 ± 1.474 |
| S-$P_{N180Q}$ | 119.63 ± 1.187 | 0.0853± 0.0139 | 94.96 ± 0.529 |
| MB-$P_{N180Q}$ | 123.67 ± 2.677 | 0.0770 ± 0.0195 | 95.22 ± 2.802 |
| eGFP | 135.95 ± 8.258 | 0.1245 ± 0.0428 | 91.79 ± 3.514 |
